# Multiepitope-based vaccine design against DiiA in *Streptococcus pneumoniae*, An immunoinformatics approach

**DOI:** 10.1101/2023.09.26.559565

**Authors:** Mariam M. Soliman, Dania Sheikhani, Jihan Nassar, Sherif Elsabbagh, Tamer M. Ibrahim

**Affiliations:** Department of Biotechnology, Faculty of Agriculture, Cairo university, Cairo, Egypt; Faculty of Pharmaceutical engineering and Biotechnology, German International University, Cairo, Egypt; Faculty of Biotechnology, Misr University for Science and Technology, Cairo, Egypt; Department of Biochemistry, Faculty of Pharmacy, University of Tübingen, Tübingen, Germany; Department of Pharmaceutical Chemistry, Faculty of Pharmacy, Kafrelsheikh University, Kafrelsheikh, 33516, Egypt

## Abstract

*Streptococcus pneumoniae* (SPN) infection has resulted in significant morbidity and mortality worldwide in children and adults. It is responsible for colonizing the human nasopharynx and can also cause diseases, including otitis media, pneumonia, bacteraemia, and meningitis. SPN is exhibiting resistance to multiple antibiotics and current vaccines have a number of limitations including poor immunogenicity and limited effectiveness against all pneumococcal serotypes. Here, we explain the design of a novel multi-epitope vaccine using Dimorphic invasion-involved protein A (DiiA) as a target protein. For designing the vaccine, the sequence of DiiA was obtained and various bioinformatics tools were employed to predict multiple CTL, HTL, B lymphocyte epitopes of DiiA. After evaluating antigenicity, allergenicity, toxicity, and immunogenicity, the most promising epitopes were chosen for constructing the vaccine, followed by an analysis of their physicochemical and immunological characteristics.The prediction, refinement, and validation of the 3D structure of the vaccine were carried out. Molecular docking, molecular dynamic simulation, and immune simulation were executed to examine the binding affinities and biological interactions at the atomic level between the vaccine and Toll-like receptor (TLR4). Vaccine translation, codon optimization were performed and expression efficiency was assessed through an in-silico cloning experiment performed to transfer into pET28a(+) plasmid vector.The obtained results proved that the vaccine maintained structural stability and possessed the capability to stimulate an efficient immune response against *S. pneumoniae* infection. The constructed vaccine has the potential for subsequent wet lab experimentation, leading to the development of an innovative vaccine.

## Introduction

Streptococcus pneumoniae (SPN), commonly known as pneumococcus, is a significant causative agent of community-acquired pneumonia, leading to a substantial number of deaths worldwide (Mazumder, Shahab et al. 2023). A variety of infections brought on by this bacterium are referred to as pneumococcal diseases, which can have serious effects on global health (Brooks and Mias 2018). Infected people have a nasopharyngeal infection that is initially asymptomatic until SPN establishes itself as a commensal flora in the upper respiratory tract (URT) (Varghese, Jayaraman et al. 2017) (Bittaye and Cash 2015). This gram-positive bacterium exhibits an elongated, round shape, is alpha-hemolytic, and possesses a capsule, while being non-motile and non-spore-forming (Mazumder, Shahab et al. 2023). Beyond pneumonia, pneumococcal infections can result in various other serious health conditions, including bronchitis, brain abscess, otitis media, septicemia, meningitis, osteomyelitis, cellulitis, pericarditis, endocarditis, conjunctivitis, peritonitis, and acute sinusitis (Mitchell and Mitchell 2010, Brooks and Mias 2018, Mazumder, Shahab et al. 2023). By interfering with immune responses and producing harmful toxins, these pathogens disrupt the normal functioning of the human body. Pneumococcal diseases can spread easily through different modes, such as person-to-person contact, indirect contact, or transmission via animal vectors like mosquitoes and ticks (Brooks and Mias 2018). Age plays a crucial role in the risk of developing pneumococcal pneumonia, with a significantly higher incidence observed in children under 2 years of age and adults over 65 years of age compared to adolescents (Ortqvist, Hedlund et al. 2005). The impact of SPN infections is particularly severe among vulnerable populations, as evidenced by the substantial number of deaths, approximately 335,000, attributed to pneumococcal diseases in children in 2015 (Mazumder, Shahab et al. 2023). Notably, regions such as South Asia and South Africa have experienced notable mortality rates, emphasizing the global burden of pneumococcal infections (Kumar, Arora et al. 2016, von Mollendorf, Tempia et al. 2017).

Understanding the role of virulence factors in bacterial colonization and pathogenesis of infection is of paramount importance for designing and developing successful peptide-based vaccines (Mitchell and Mitchell 2010, Rafi, Al-Khafaji et al. 2023). These virulence factors may help SPN in spreading in the host cell, escaping the immune defense system, and promoting disease progression. Many in-silico studies evaluated these virulence proteins as vaccine candidates for epitope-based vaccine design against SPN (Mazumder, Shahab et al. 2023). SPN expresses several virulence factors, such as Polysaccharide capsule, the pore□forming toxin, pneumolysin (PLY), LPXTG motif-anchored surface proteins containing hyaluronidase, neuraminidase, and serine protease PrtA, pili, choline-binding proteins (CBPs),, pneumococcal surface proteins (PspA and PspC), pneumococcal histidine triad (Pht) proteins (PhtD and PhtE), and the lipoprotein components of iron uptake ATP-binding cassette (ABC) transporters (PiuA, PiaA, and PsaA) (Subramanian, Henriques-Normark et al. 2019, Mazumder, Shahab et al. 2023).

Dimorphic invasion-involved protein A (DiiA) is a pneumococcal surface protein encoded by the sp_1992 locus of the TIGR4 strain. DiiA has two major allelic variants of different lengths carrying either one (R2) or two (R1 and R2) amino terminal repeats (Martin-Galiano, Escolano-Martinez et al. 2021). It has a typical topology of adhesins, which consists of a conserved C-terminal LPxTG cell wall anchor, a non-repeat unstructured coiled-coil stalk, and repeated motifs that differ in number between alleles. The clinical isolates showed a correlation between the allele and invasiveness. Invasive pneumococcal disease (IPD) is related to the long allele, and the noninvasive disease (non-IPD) is related to the short allele (Escolano-Martínez, Domenech et al. 2016, Martin-Galiano, Escolano-Martinez et al. 2021).

DiiA is a virulence factor contributing to two stages of the IPD process. The first stage is the internalization through the lungs and infection establishment which is attributed to the cooperation of R2 with R1. The second stage is the dissemination to the systemic circulation, primarily due to the non-repeat unstructured coiled-coil stalk (Escolano-Martínez, Domenech et al. 2016). The unstructured section of DiiA has a crucial role in bacterial proliferation in blood during pneumococcal sepsis by avoiding fast pneumococcal clearance, besides, interacting with collagen and lactoferrin, two fundamental proteins of the innate immunity. In addition, DiiA-NR mutants exhibited a significant decrease in bacterial replication in the upper and lower respiratory tracts (Escolano-Martínez, Domenech et al. 2016, Martin-Galiano, Escolano-Martinez et al. 2021).Because of its limited allelic variability, immunogenicity, high global B-cell antigenic characteristics, non-toxicity, and non-allergenicity we considered DiiA as a suitable target antigen of SPN for multi-epitope vaccine design.

Vaccination continues to be the preeminent approach in the prevention and management of infectious diseases. A multitude of established vaccines, primarily those of the live attenuated, inactivated (killed) microorganisms and subunit varieties, result in the efficacious elicitation of safeguarding immune responses, primarily through the mediation of antibody-based responses against pathogens. Nonetheless, it has come to our attention that a vast array of exceedingly hazardous pathogens cannot be regulated through traditional vaccination techniques (Soleymani, Tavassoli et al. 2021). The development and production of vaccines are associated with significant costs and time investments, often requiring several years to achieve(Ribas□Aparicio Rosa, Castelán□Vega Juan et al. 2017). Recombinant vaccines, such as conjugated, subunit, and toxoid vaccines, may cause severe toxicity instead of the intended immune response(Mazumder, Shahab et al. 2023).

There are now two types of pneumococcal vaccines on the market: protein-conjugated polysaccharide vaccines (PCV) and polysaccharide vaccines (PPV) based on capsular polysaccharides of at least 92 distinct serotypes (Dorosti, Eslami et al. 2019). Four distinct pneumococcal vaccines (PPSV23, PCV7, PCV10, and PCV13) have been made available to the public since 1983; each has its own drawbacks and advantages(Wantuch and Avci 2018). PCVs like PCV7, PCV10, and PCV13—which cover 7, 10, and 13 serotypes, respectively— highly protect newborns, PPVs like PPV23 do not effectively stimulate protective immunity in children under the age of two. However, PCVs have some drawbacks, including complicated manufacturing, expensive production, the need for refrigeration, and multiple injections. A promising replacement for the current capsular antigen vaccines is the use of epitope-based vaccinations, which may contain various combinations of conserved virulence proteins (Wantuch and Avci 2018). Numerous strategies have been implemented with the aim of mitigating these challenges, primarily concentrating on the identification of suitable antigens or antigenic structures, carriers, and adjuvants to streamline the development process and reduce associated costs (Plotkin 2014).

Bioinformatics techniques can help scientists in a variety of biological domains anticipate possibly immunoprotective epitopes in design candidate vaccines, the design of immunodiagnostic modalities, and the production of antibodies (Kazi, Chuah et al. 2018, Shafaghi, Bahadori et al. 2023). The immuno-bioinformatics technologies really have benefits over traditional vaccination techniques, such as quicker results in just one to two years and reduced expenses (Soleymani, Tavassoli et al. 2021, Shafaghi, Bahadori et al. 2023). By examining the pathogen’s entire genome to identify possible antigens, reverse vaccinology aids in epitope mapping and predicts monovalent peptide vaccines, leading to the development of a potential pneumococcal vaccination using a novel strategy (Oli, Obialor et al. 2020, Sunita, Sajid et al. 2020).

The initial stage employed in reverse vaccinology entails dry lab analysis to determine whether the vaccine design is novel and effective in inducing both humoral and cellular immunities essential for protection against S. pneumoniae (Kazi, Chuah et al. 2018, Dorosti, Eslami et al. 2019). Computational tools and online databases are utilized to select a gene of interest. Further analysis of the chosen genomes is conducted to identify potential immunogenic epitopes that can stimulate an immune response in B-cells or T-cells. Once the antigenic candidates have been selected, they are then subjected to the second phase (wet lab) for validation. In this latter phase, antigenic proteins or peptides of interest are produced or synthesized (Kazi, Chuah et al. 2018). In silico vaccine construction is hindered by unreliable analysis that requires verification through wet laboratory experiments. Computational databases have limitations that may lead to spurious outcomes. Therefore, in vitro testing and clinical trials in larger animals are necessary to ensure the efficacy of vaccines produced by in silico experimentation(Mazumder, Shahab et al. 2023).

The aim of this study is to predict potential epitopes (T and B cells epitopes) from the DiiA protein. Identified epitopes will be assessed for their antigenicity, immunogenicity, and allergenicity and will be further joined with suitable adjuvants and linkers to construct a multi epitope vaccine. Molecular docking and molecular dynamics simulation will be performed to study the interaction between the vaccine and toll-like receptor and analyze the stability of the docked complex. Finally, the immune response elicited by the constructed vaccine will be analyzed by in-silico immune simulation.

## Materials and Methods

### Protein sequence retrieval and analysis

Sequence of DiiA (SP_1992) from *Streptococcus pneumoniae* strain ATCC BAA-334 / TIGR4 was retrieved from uniprot database (ID: A0A0H2URQ5). VaxiJen v2.0 (http://www.ddg-pharmfac.net/vaxijen/VaxiJen/VaxiJen.html) (Doytchinova and Flower 2007) and AllerTOP 2.0 (https://www.ddg-pharmfac.net/AllerTOP/) (Dimitrov, Bangov et al. 2014) servers were used to determine the antigenicity and allergenicity of the retrieved sequence respectively. A value of 0.4 was used as a threshold for the VaxiJen server. A sequence similarity search of the DiiA protein against human proteome was conducted using NCBI BlastP to avoid cross reactivity. Physicochemical properties of the target protein were determined using the ProtParam tool of ExPasy server (https://web.expasy.org/protparam/) (Wilkins, Gasteiger et al. 1999).

### Prediction of CTL epitopes

NetMHC v4.0 (https://services.healthtech.dtu.dk/services/NetMHC-4.0/) was used to predict the cytotoxic T lymphocyte epitopes of the DiiA protein sequence (Andreatta and Nielsen 2016). The peptide length was set to 9-mers, and the thresholds for strong binders and weak binders were set to 0.5 and 2 respectively. All the alleles of the HLA supertype representative (HLA-A0101, HLA-A0201, HLA-A0301, HLA-A2402, HLA-A2601, HLA-B0702, HLA-B0801, HLA-B2705, HLA-B3901, HLA-B4001, HLA-B5801 and HLA-B1501) were selected. The resulting epitopes were further evaluated for their antigenicity, allergenicity, toxicity, and immunogenicity using VaxiJen v2.0, AllerTop v2.0, ToxinPred (https://webs.iiitd.edu.in/raghava/toxinpred/multi_submit.php) (Gupta, Kapoor et al. 2013), and IEDB (http://tools.iedb.org/immunogenicity/) (Calis, Maybeno et al. 2013) with default parameters, respectively.

### Prediction of HTL epitopes

HTL (Helper T lymphocyte) epitopes were identified for potential vaccine development using the NetMHCII-2.3 server https://services.healthtech.dtu.dk/services/NetMHCII-2.3/) (Jensen, Andreatta et al. 2018). During the HTL prediction, the NN-align approach and the entire set of alleles were chosen.The MHC class II alleles were predicted using the human alleles DR, DP, and DQ. Based on their assigned percentile ranks, the epitope sequences were classified as strong binders (SBs), weak/intermediate binders (WBs), and non-binders (NBs). To identify the most suitable epitopes for vaccine preparation, a comprehensive analysis was conducted. This analysis included the assessment of antigenicity, allergenicity, and toxicity using VaxiJen v2.0, AllerTop v2.0, and ToxinPred tools, respectively. Furthermore, the IFNepitope server (https://webs.iiitd.edu.in/raghava/ifnepitope) was utilized to analyze the potential IFN-γ response (Dhanda, Vir et al. 2013).

### Linear B lymphocyte epitopes prediction

B cell epitopes are essential for humoral or antibody-mediated immunity, so they are considered an important component in vaccine construction. The ABCpred server (https://webs.iiitd.edu.in/raghava/abcpred/index.html) was used to predict linear B cell epitope (LBL) regions in the sequence of DiiA using an artificial neural network with a threshold value of 0.51, and 16-mers were selected as the preferred window length (Saha and Raghava 2006). Then we evaluated the predicted epitopes’ antigenicity, allergenicity, and toxicity indexes using VaxiJen v2.0, AllerTOP 2.0, and ToxinPred servers, respectively.

### Multi-epitope vaccine construction

To stimulate both innate and adaptive immune responses, a multi-epitope vaccine was constructed using the predicted CTL, HTL and LBL epitopes from DiiA protein. For a stronger immune response, 50S ribosomal protein L7/L12 (Uniprot ID: P9WHE3) was used as an adjuvant. It was linked at the N-terminal to CTL epitopes using the “EAAAK” linker followed by HTL and LBL epitopes. CTL and HTL epitopes were linked by “AAY” and “GPGPG” linkers respectively whereas “KK” linker was utilized to link LBL epitopes.

### Physicochemical properties and immunological analysis

To determine the physicochemical properties of the vaccine, the ExPasy ProtParam server was used. Various physical and chemical properties of the vaccine can be identified, such as the number and composition of amino acid residues, the number of positively and negatively charged residues, the molecular weight, the theoretical pI, the estimated half-life in different cell types, the aliphatic and instability index, and the grand average of hydropathicity (GRAVY). In addition, the assessment of the vaccine’s immunological characteristics involved the utilization of VaxiJen v2.0, AllerTop. SOLpro server (http://scratch.proteomics.ics.uci.edu/) (Cheng, Randall et al. 2005) was used to assess the vaccine solubility.

### Population coverage and conservancy analysis

Population coverage of the chosen epitopes was evaluated by the IEDB population coverage tool (http://tools.iedb.org/population/) (Bui, Sidney et al. 2006). Each epitope coverage was assessed against MHC class I and II binding alleles using the combined option. The degree of conservancy of the epitopes was analyzed using the IEDB conservancy analysis tool (http://tools.iedb.org/conservancy/).

### Vaccine 3D structure prediction, refinement and validation

The three-dimensional structure of the constructed vaccine was predicted using the robetta server (https://robetta.bakerlab.org/) (Kim, Chivian et al. 2004). RoseTTAFold, a deep learning-based modeling method was selected. GalaxyRefine server (https://galaxy.seoklab.org/cgi-bin/submit.cgi?type=REFINE) was utilized to further refine the predicted model (Ko, Park et al. 2012). Structural validation of the refined model was performed using ERRAT (Colovos and Yeates 1993) and PROCHEK (R. A. Laskowski 1993) from the SAVES server (https://saves.mbi.ucla.edu/).

### Conformational-B cell epitope prediction

Discontinuous B-cell epitopes were predicted in the modeled vaccine 3D structure using Ellipro server (http://tools.iedb.org/ellipro/)(Ponomarenko, Bui et al. 2008).

### Molecular docking

Molecular docking was performed to predict the interaction between the multi-epitope vaccine and TLR4. A protein-protein docking was done using ClusPro 2.0 (https://cluspro.bu.edu/) while keeping all the docking parameters as default (Kozakov, Hall et al. 2017). The structure of TLR4 (PDB ID: 3FXI) was retrieved from the RCSB protein database (PDB) and then prepared for docking by removing associated ligands and water molecules. The docked complex was visualized by PyMOL software. Finally, the intermolecular interactions between the vaccine and the receptor were investigated with the PDBsum tool (http://www.ebi.ac.uk/thornton-srv/databases/pdbsum/Generate.html) (Laskowski, Jablonska et al. 2018).

### Molecular dynamics (MD) simulation

Structural dynamics of the TLR4-vaccine complex was investigated using the iMODS server (http://imods.chaconlab.org/) (Lopez-Blanco, Aliaga et al. 2014). It uses normal mode analysis (NMA) to calculate internal coordinates of the protein for stability assessment and the results are shown as deformability plot, B-factor value, covariance matrix, elastic network map and eigenvalue.

### Immune simulation

Computational immune simulation was performed using the C-IMMSIM server (https://kraken.iac.rm.cnr.it/C-IMMSIM/) (Rapin, Lund et al. 2010). The C-IMMSIM server uses a position-specific scoring matrix (PSSM), which can comprehend an immune response that is produced because of vaccination dosage over time. The designed epitope construct in the FASTA format is submitted to the server. Three doses with default parameters were injected. A total of 1050 simulation steps (which cover the time period of a year) were chosen. The time steps were set at 1, 84, and 170 (time step 1 is injection at time = 0, and each time step is 8 h). Two MHC class I alleles (B0702 and B3901) and two MHC class II alleles (DRB1_0403 and DRB1_0404) were selected. This was carried out to simulate and contrast the host’s immune system after vaccination.

### Codon adaptation and in silico cloning

Java Codon Adaptation Tool (JCat) server (http://www.jcat.de/) (Grote, Hiller et al. 2005) was used for the reverse translation and optimization of the DNA sequence to adapt its codon to most sequenced prokaryotic organisms *E. coli* strain (K12). GC content and a score for codon adaptation index (CAI) were measured for the adapted and non-adapted sequences. XhoI and NdeI sites were attached to 5’ and 3’ ends of the nucleic acid sequence and the codon-optimized (adapted) DNA sequence of the vaccine was introduced into the *E. coli* pET28a(+) vector using the SnapGene software through a cloning procedure.

## Results

### Protein sequence retrieval and analysis

Prior to various epitopes prediction, DiiA sequence was obtained and analyzed for its antigenicity and allergenicity. The protein was predicted to be highly antigenic with VaxiJen score of 0.9798. AllerTOP server predicted DiiA to be probable non-allergen. In addition, the protein was found to be non-homologous to the human proteome. These results indicated the DiiA protein as a suitable target for vaccine construction. The physicochemical properties of the protein are listed in Supplementary Table 1.

**Table 1.**
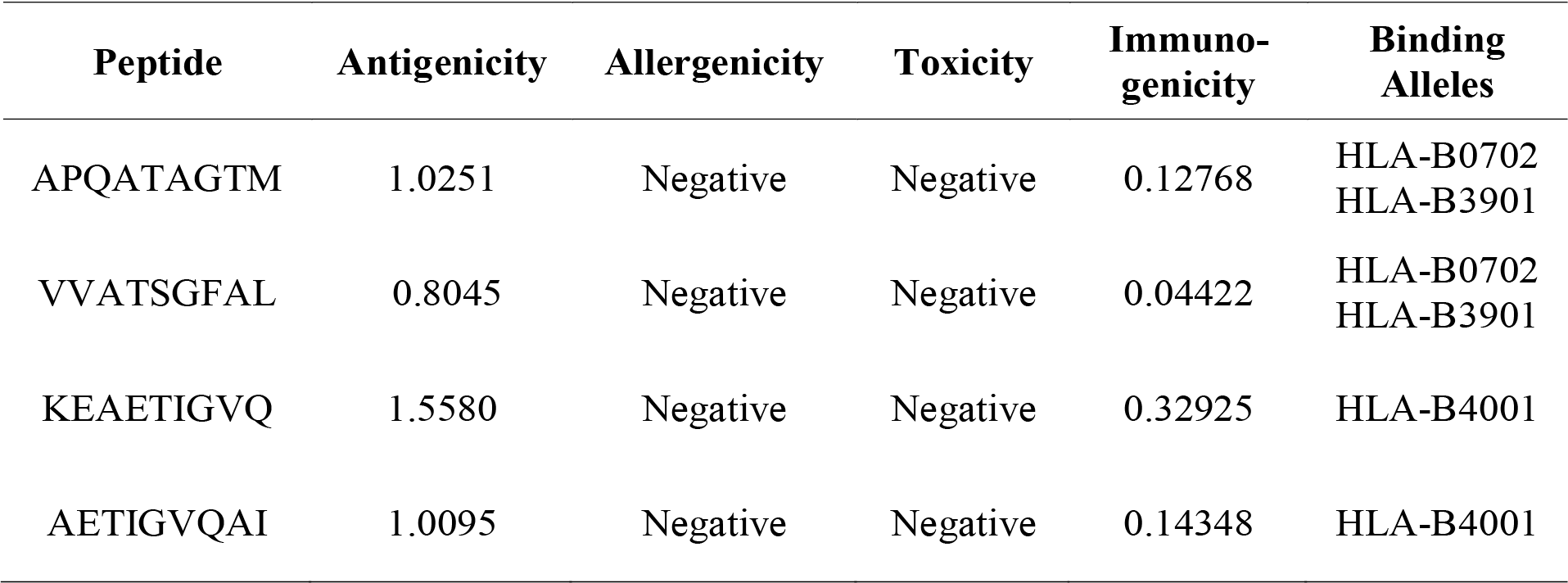
CTL epitopes identified in DiiA sequence using NetMHC v4.0 server.

### Prediction of CTL epitopes

CTL epitopes, along with HTL and LBL, are required for the construction of a multi-epitope vaccine that provides long-lasting immunity [1,3]. Therefore, CTL epitopes of DiiA protein were predicted using the NetMHC v4.0 server. Several epitopes were obtained, which were further filtered based on their antigenicity, immunogenicity, toxicity, and allergenicity to eventually obtain the best four epitopes to be used for vaccine construction. The selected four epitopes are provided in Table 1 with their antigenicity and immunogenicity scores, as well as the MHC I alleles they can bind to.

### Prediction of HTL epitopes

HTL epitope prediction is essential when formulating a multi-epitope vaccine. T-helper cells (CD4+) assume a pivotal role in regulating and optimizing the immune response against pathogens. Besides their regulatory function, T-helper cells also stimulate innate immune cells, B lymphocytes, and cytotoxic T cells, which together form a comprehensive and effective immune defense mechanism (Rapin, Lund et al. 2010, Mazumder, Shahab et al. 2023). Among the epitopes predicted by NetMHCII-2.3 listed in Supplementary Table 2, 4 epitopes were selected for vaccine construction. They were found to be highly antigenic, non-allergenic, non-toxin, immunogenic and can interact with many alleles (Table 2).

### Prediction of LBL epitopes

B-cell epitopes are surface-accessible clusters of amino acids that are identified by secreted antibodies or B-cell receptors, function as cells that present antigens to the immune system and cause an immunological response. A crucial first step in the development of vaccines and therapeutic antibodies based on epitopes is the identification of B-cell epitopes. In our study, the ABCpred service generated 16 peptide sequences for B cell epitopes (Supplementary Table 3). We looked at the toxicity, allergenicity, and antigenicity of the chosen epitopes. We analyzed the 16 peptide sequences and found 10 peptides to be antigenic, non-allergenic, and nontoxic, as listed in Table 3. The top four epitopes among them with the best antigenicity, allergenicity, toxicity, and allergenicity were chosen for the final vaccine construction.

**Table 2.** HTL epitopes predicted using NetMHCII-2.3 server.

| Peptide | Antigenicity | Allergenicity | Toxicity | IFN-γ score | Interacting alleles |
| --- | --- | --- | --- | --- | --- |
| ASLGLVVATSGF<br>ALL | 0.4453 | Negative | Negative | 0.54893638 | DRB1_0402 |
|  |  |  |  |  | DRB1_0403 |
|  |  |  |  |  | DRB1_0404 |
|  |  |  |  |  | DRB1_0701 |
|  |  |  |  |  | DRB1_0901 |
|  |  |  |  |  | DRB3_0301 |
|  |  |  |  |  | HLA-DPA10103- |
|  |  |  |  |  | DPB10401 |
|  |  |  |  |  | HLA-DPA10103- |
|  |  |  |  |  | DPB10201 |
|  |  |  |  |  | HLA- |
|  |  |  |  |  | DQA10501- |
|  |  |  |  |  | DQB10302 |
|  |  |  |  |  | HLA- |
|  |  |  |  |  | DQA10501- |
|  |  |  |  |  | DQB10301 |
|  |  |  |  |  | HLA- |
|  |  |  |  |  | DQA10501- |
|  |  |  |  |  | DQB10303 |
|  |  |  |  |  | HLA- |
|  |  |  |  |  | DQA10201- |
|  |  |  |  |  | DQB10303 |
|  |  |  |  |  | HLA- |
|  |  |  |  |  | DQA10103- |
|  |  |  |  |  | DQB10603 |
|  |  |  |  |  | HLA- |
|  |  |  |  |  | DQA10102- |
|  |  |  |  |  | DQB10602 |
|  |  |  |  |  | HLA- |
|  |  |  |  |  | DQA10102- |
|  |  |  |  |  | DQB10501 |
|  |  |  |  |  | HLA- |
|  |  |  |  |  | DQA10201- |
|  |  |  |  |  | DQB10301 |
|  |  |  |  |  | HLA- |
|  |  |  |  |  | DQA10301- |
|  |  |  |  |  | DQB10301 |
| LASLGLVVATSG<br>FAL | 0.4650 | Negative | Negative | 0.36598918 | DRB1_0403 |
|  |  |  |  |  | DRB1_0402 |
|  |  |  |  |  | DRB1_0901 |
|  |  |  |  |  | HLA- |
|  |  |  |  |  | DQA10201- |
|  |  |  |  |  | DQB10301 |
|  |  |  |  |  | HLA- |
|  |  |  |  |  | DQA10501- |
|  |  |  |  |  | DQB10303<br>HLA-DQA10102-DQB10501<br>HLA-DQA10102-DQB10602<br>HLA-DQA10103-DQB10603<br>HLA-DQA10201-DQB10303<br>HLA-DQA10301-DQB10301<br>HLA-DQA10501-DQB10301<br>HLA-DQA10501-DQB10302 |
| KTTSAPQATAGT<br>MQD | 1.0114 | Negative | Negative | 0.20456662 | HLA-DQA10501-DQB10302<br>HLA-DQA10501-DQB10301<br>HLA-DQA10501-DQB10303<br>HLA-DQA10201-DQB10301<br>HLA-DQA10201-DQB10303 |
| APNAAPKTTSAP<br>QAT | 0.8274 | Negative | Negative | 0.54291746 | HLA-DQA10201-DQB10301<br>HLA-DQA10501-DQB10301<br>HLA-DQA10501-DQB10303 |

**Table 3.**
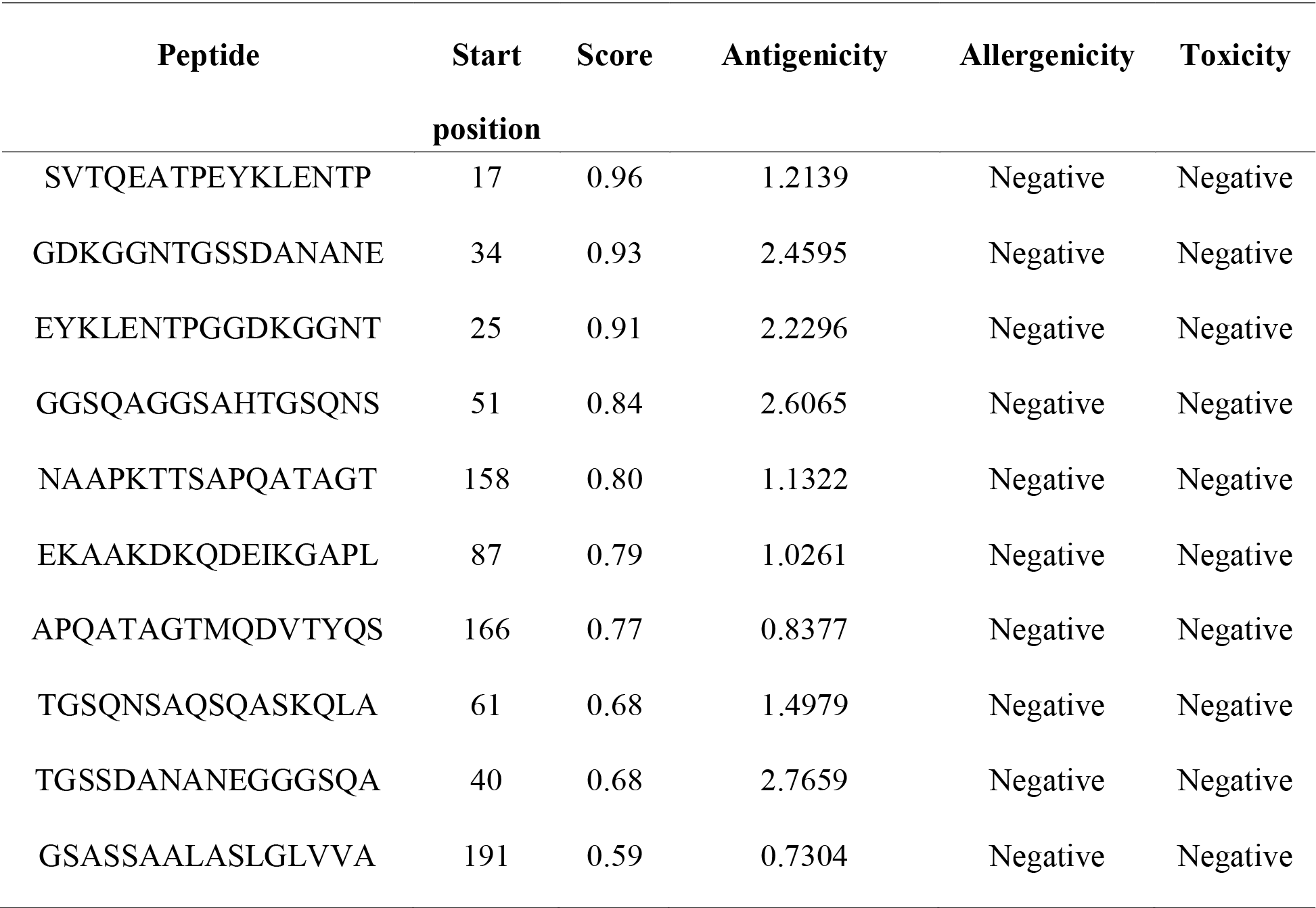
Predicted antigenic B cell epitopes found in the DiiA protein.

### Multi-epitope vaccine construction

Following various epitopes prediction and screening, a multi-epitope vaccine was constructed. A total of 12 epitopes were used comprising 4 CTL, 4 HTL, and 4 LBL epitopes. 50S ribosomal protein being selected as an adjuvant was attached to CTL epitopes using EAAAK linker. The CTL epitopes were linked together by the “AAY” linker which enhances vaccine immunogenicity and affects epitope presentation capacity. “GPGPG” and “KK” which were used to link HTL, and B cell epitopes respectively are known to influence immune processing and immunogenic activity of the vaccine candidate (Nain, Abdulla et al. 2020). The final vaccine construct composed of 332 amino acids is shown in Figure 1.

**Figure 1.**
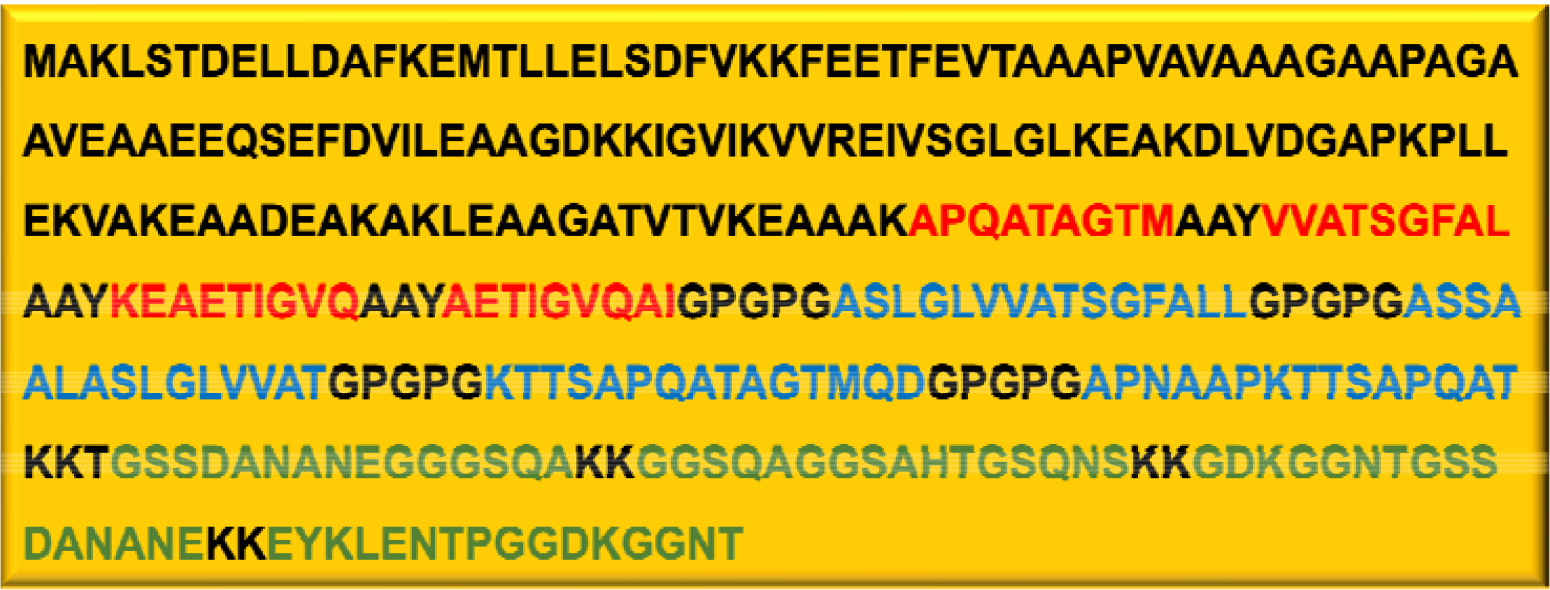
Sequence of the designed vaccine. The CTL, HTL and LBL epitopes are depicted in red, blue, and green colors respectively.

### Physicochemical properties and immunogenicity analysis

The obtained results from the ExPasy ProtParam server revealed that the vaccine construct consists of a total of 332 amino acids with a molecular weight of 32364.16 Da. The theoretical isoelectric point value (pI) was calculated to be 5.06, indicating the acidic nature of the vaccine. The half-life was estimated to be 30 hours in mammalian reticulocytes, >20 hours in yeast, and >10 hours in Escherichia coli. The instability index, aliphatic index, and grand average of hydropathicity (GRAVY index) were calculated as 19.93, 73.77, and -0.161, respectively, indicating a stable vaccine with high thermostability and hydrophilic properties. VaxiJen v2.0 predicted it as antigenic with as score of 1.0518. AllerTop v2.0 predicted the vaccine to be non-allergenic, making it appropriate for vaccine development. SOLpro also indicated that the sequence had a high probability of solubility upon overexpression in *E*.*coli*, scoring 0.663710.

### Population coverage and conservancy analysis

HLA allele distribution should be considered when developing a successful vaccine. For this, the population coverage of the designed vaccine was evaluated using the IEDB tool. Analysis showed that the selected epitopes in our vaccine would cover 99.02% of the world population (Table 4). It is worth noting that all world regions showed very high population coverage except for south africa which had a coverage of 8.81%. The conservancy analysis showed that the epitopes used in the vaccine construct are 100% conserved among the related protein sequences.

**Table 4.**
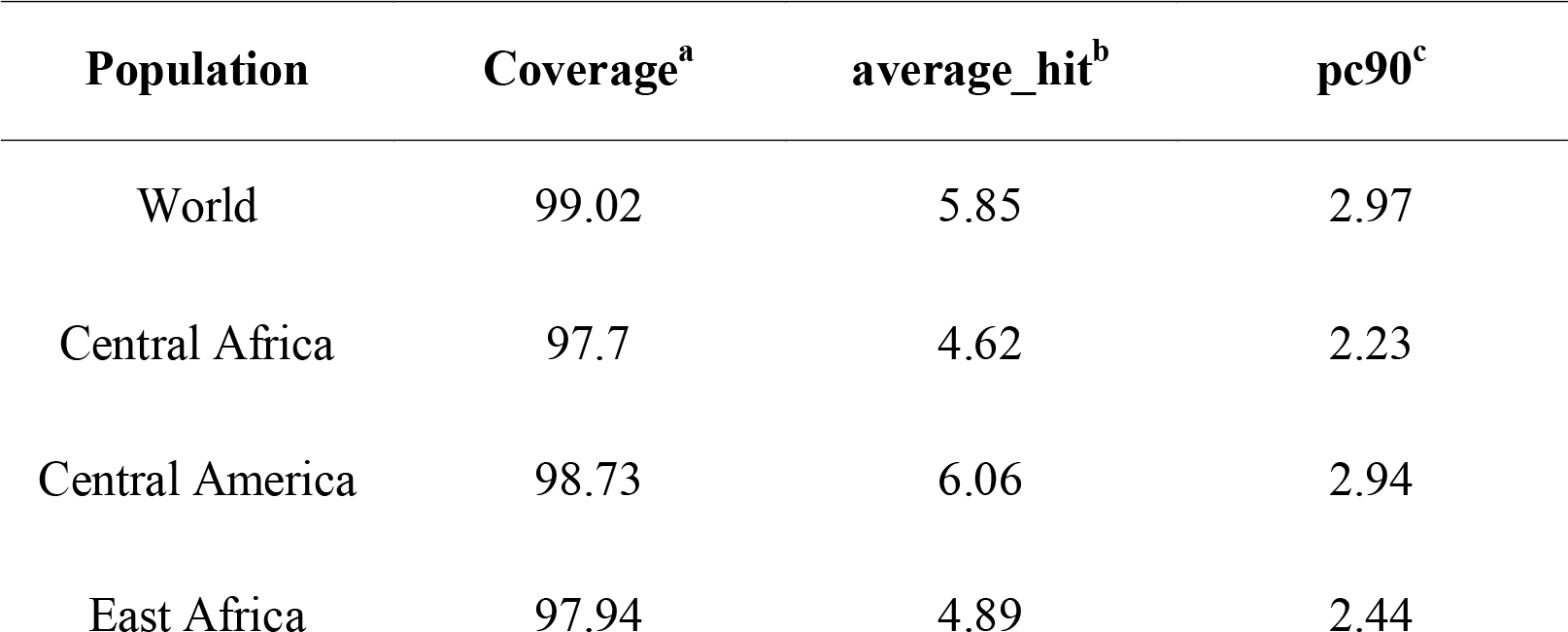
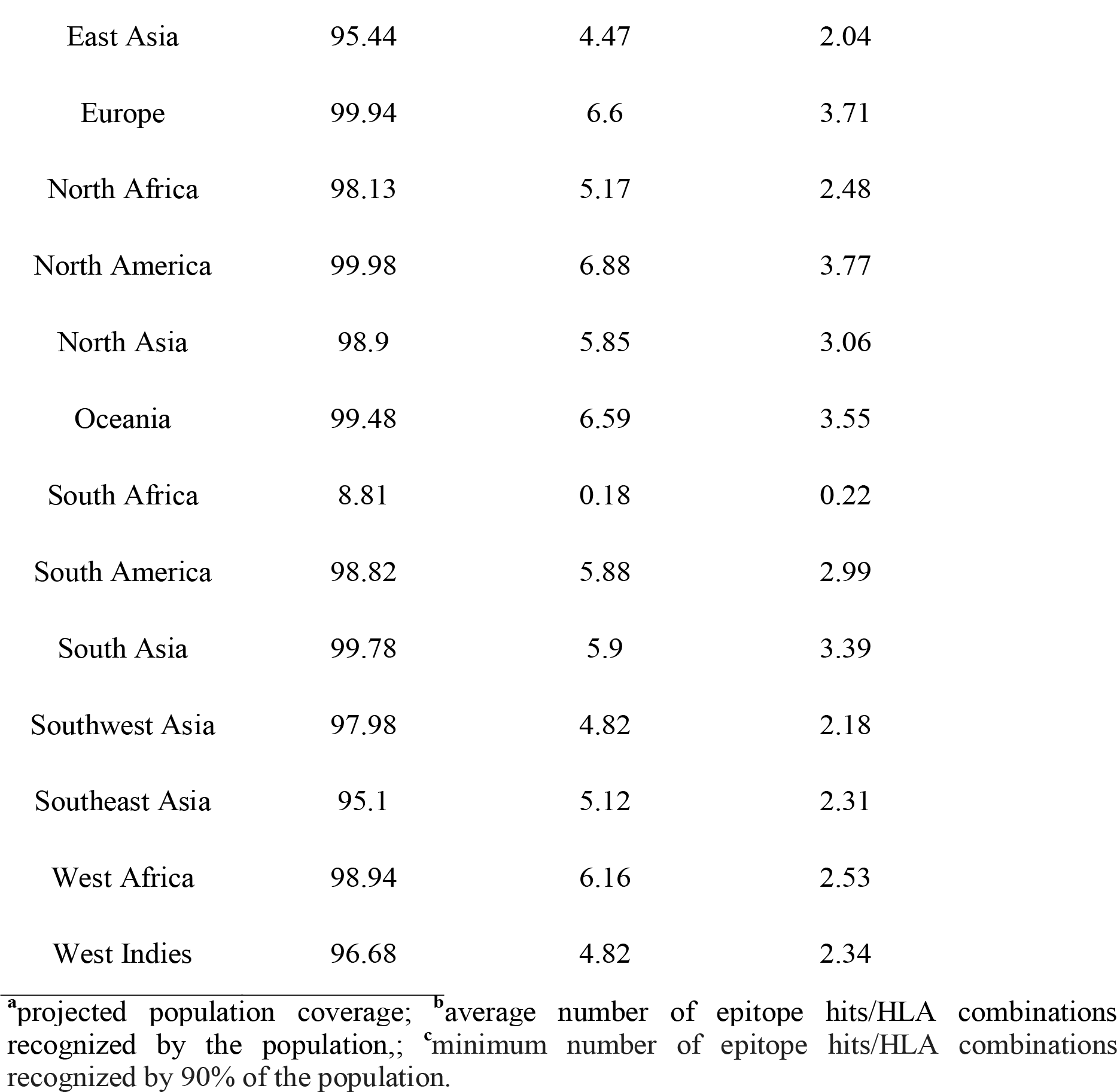
Population coverage of selected epitopes.

### Vaccine 3D structure prediction, refinement, and validation

The designed multi-epitope vaccine structure was modeled by the Robetta server using the deep learning approach. 5 models were generated, and the first model was subjected to refinement by the GalaxyRefine server. Model number 2 was selected, among the five refined models for further analysis (Figure 2A). It showed a GDT-HA score of 0.9744, MolProbity score of 1.86, clash score of 12.8, poor rotamers of 0.0 and 96.4 rama favored score. Both initial and refined models were validated by the SAVES server. The overall ERRAT quality factor of the initial model was 95.37 which decreased to 93.38 in the refined model (Figure 2B). The Ramachandran plot generated by PROCHECK displayed that for the initial model, 87.3%, 12% and 0.7% of the residues were in the favored, allowed, and disallowed regions respectively while for the refined model, 94%, 5.3% and 0.7% were in the favored, allowed, and disallowed regions respectively (Figure 2C). This indicated that the refined model had a good quality and can be used for next analysis.

**Figure 2.**
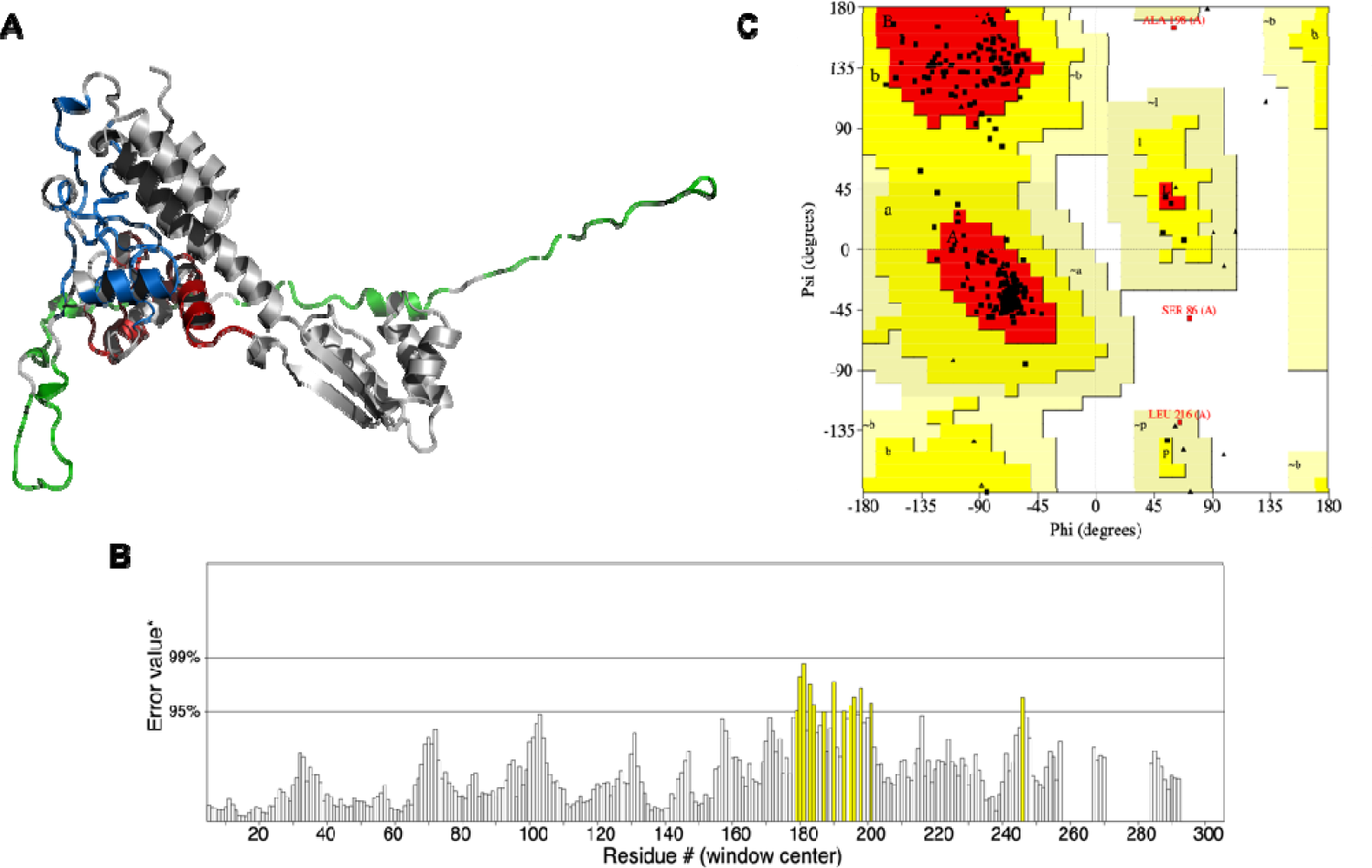
Vaccine tertiary structure prediction and validation. **(A)** Vaccine 3D model predicted and refined by Robetta and GalaxyRefine servers respectively. CTL, HTL and LBL are shown in red, blue and green colors respectively. PyMOL was used for visualization. **(B)** ERRAT plot of refined model. **(C)** Ramachandran plot generated by PROCHECK.

### Conformational-B cell epitope prediction

It has been estimated that more than 90% of the B-cell epitopes are discontinuous. They consist of residues that are far apart and are brought together into proximity through protein folding. So, it is crucial to identify them in the vaccine structure. Prediction with Ellipro server gave rise to eight conformational epitopes as depicted in Supplementary Table 4. The 3D structure of the predicted epitopes in the designed vaccine is shown in Figure 3.

**Figure 3.**
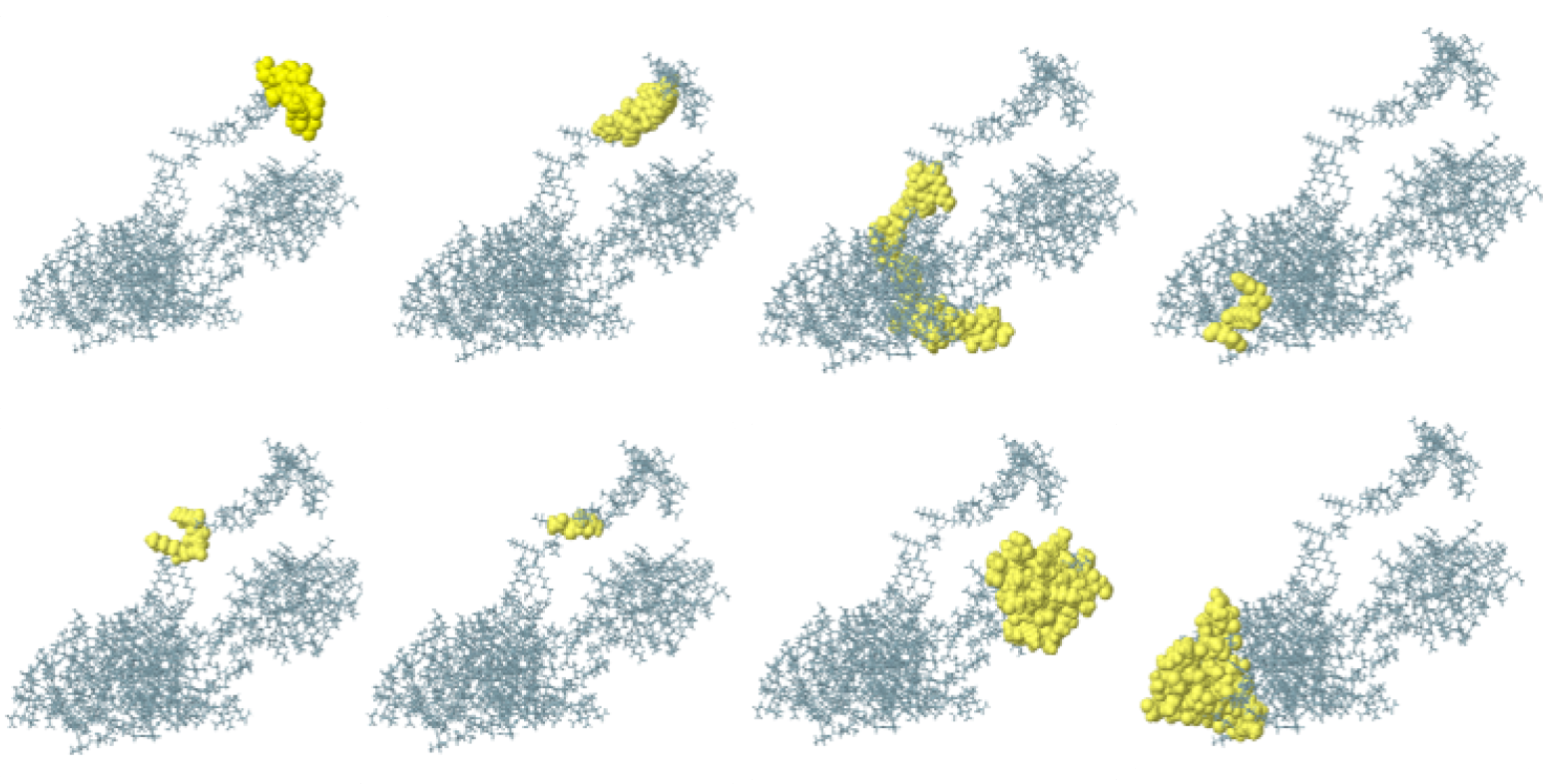
Conformational B-epitopes predicted by ellipro server. The epitopes are depicted in yellow color.

### Molecular docking

A receptor-vaccine interaction is required for a stable immune response. Hence, the binding affinities between the refined vaccine and the receptor were predicted by performing protein-protein docking using ClusPro 2.0, resulting in a total of 30 models with different energies. Among these models, model number 0 was selected as the best docked complex with a center and lowest energy weighted score of -734.6 kcal/mol. This model was selected because it contains the highest number of members in its cluster. PyMOL was used to visualize the docking interaction, shown in Figure 4A. The PDBsum server was used to reveal the interface residues and intermolecular interactions involved in the vaccine-TLR4 complex (Figure 4B, 4C). The results showed that 27 residues in the TLR4 interface covering a surface area of 956 Å2 interacted with 21 residues in the vaccine interface covering a surface area of 1034 Å2. A total of 21 hydrogen bonds, 3 salt bridges, and 199 non-bonded interactions were formed between the vaccine and the receptor. The hydrogen bonds were as follow: THR175-SER288, THR175-ALA289, THR175-HIS290,THR175-HIS290,GLU178-GLY287,LEU203-THR291,ASN205-ALA285, ASN205-GLN284, ARG227-GLN284, ARG227-GLN284, GLU254-LYS279, HIS256-SER283, ILE285-SER276, ILE285-SER276, ILE285-SER276, GLU286-LYS280, GLU286-LYS280, GLU287-LYS280, ASN309-SER276, ASN309-GLU272, and SER311-LYS280 at a distance of 2.84 Å, 3.33 Å, 2.88 Å, 2.79 Å, 3.24 Å, 3.34 Å, 3.23 Å, 2.91 Å, 2.70 Å, 2.78 Å, 2.74 Å, 2.82 Å, 2.68 Å, 3.33 Å, 2.84 Å, 2.83 Å, 2.70 Å, 2.65 Å, 2.84 Å, 2.89 Å, and 2.53 Å, respectively. Only three salt bridges were formed between GLU254-LYS279, GLU286-LYS280, and GLU287-LYS280 at a distance of 2.74 Å, 2.70 Å, and 2.65 Å, respectively.

**Figure 4.**
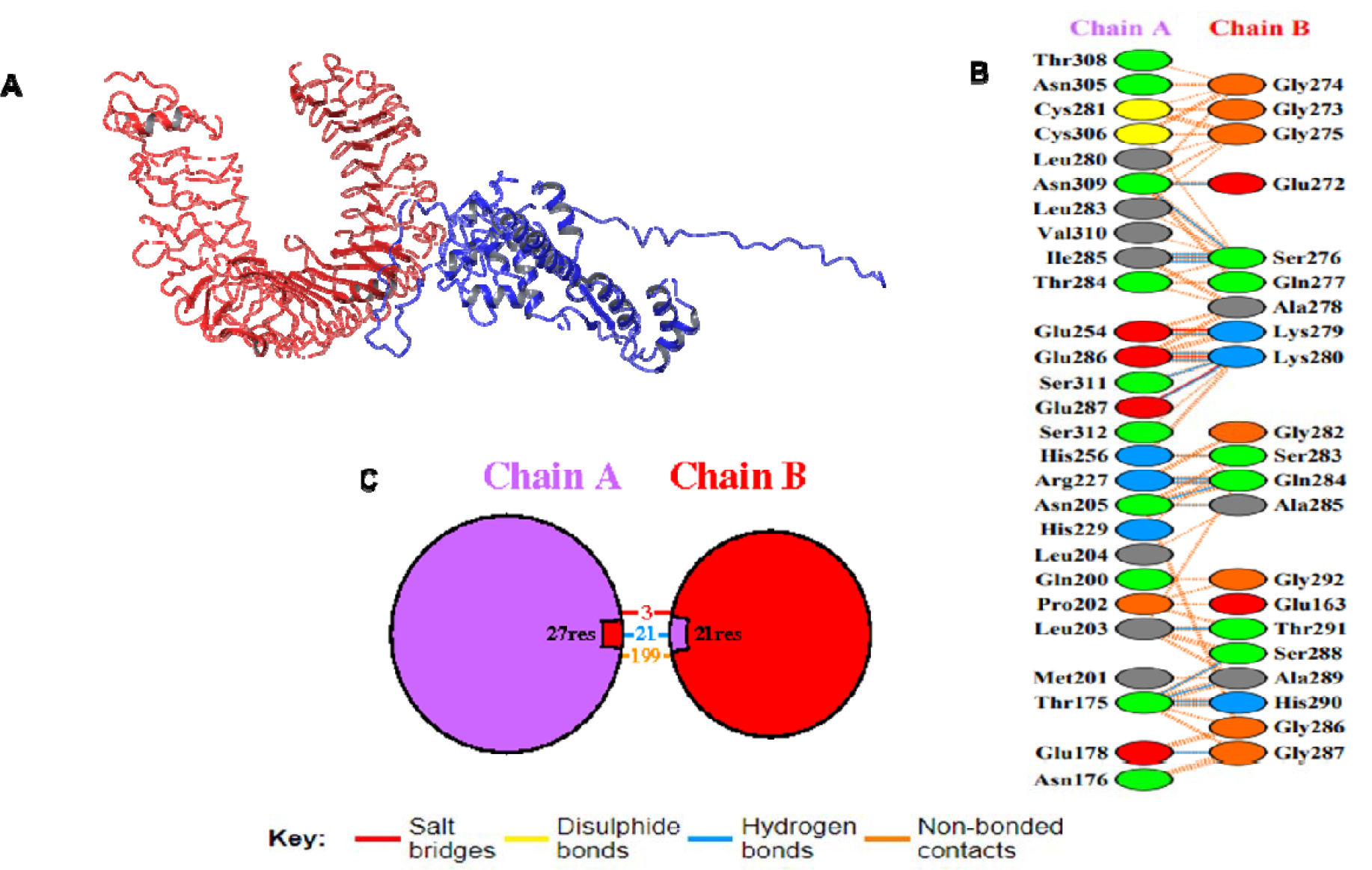
Molecular docking and interaction pattern of the vaccine construct and receptor. **(A)** Cartoon representation of the vaccine construct (blue) and TLR4 (red) docked complex illustrated with PyMOL. **(B)** Graphical illustration of the interacting residues. A total of 27 residues in the TLR4 interacted with 21 residues in the vaccine. A number of 3 salt bridges (red), 21 hydrogen bonds (blue), and 199 non-bonded interactions (orange) were formed. **(C)** Schematic diagram of interactions between TLR4 (chain A) and the vaccine (chain B).

### Molecular dynamics (MD) simulation

Stability and flexibility of the vaccine-receptor complex was evaluated using NMA employed in the iMODS server. The deformability plot showed no significant distortions except in the region between residues 650-750 and the last region in the vaccine (Figure 5A). This is of no surprise given the disordered structure adopted by this region implying high flexibility. This is confirmed by the B-factor analysis shown in Figure 5B showing almost no fluctuations except for the disordered region. The eigenvalue of the complex was 6.51e^-07^ and the variance analysis showed the stiffness of the complex (Figure 5C-D). Pairing between residues was studied using the covariance analysis where red, blue and while colors represented the correlated, anticorrelated and uncorrelated motions respectively (Figure 5E). The elastic net showed which pairs of atoms were connected by springs where each dot represented one spring. Dots were colored based on their stiffness where darker grays indicated stiffer springs (Figure 5F).

**Figure 5.**
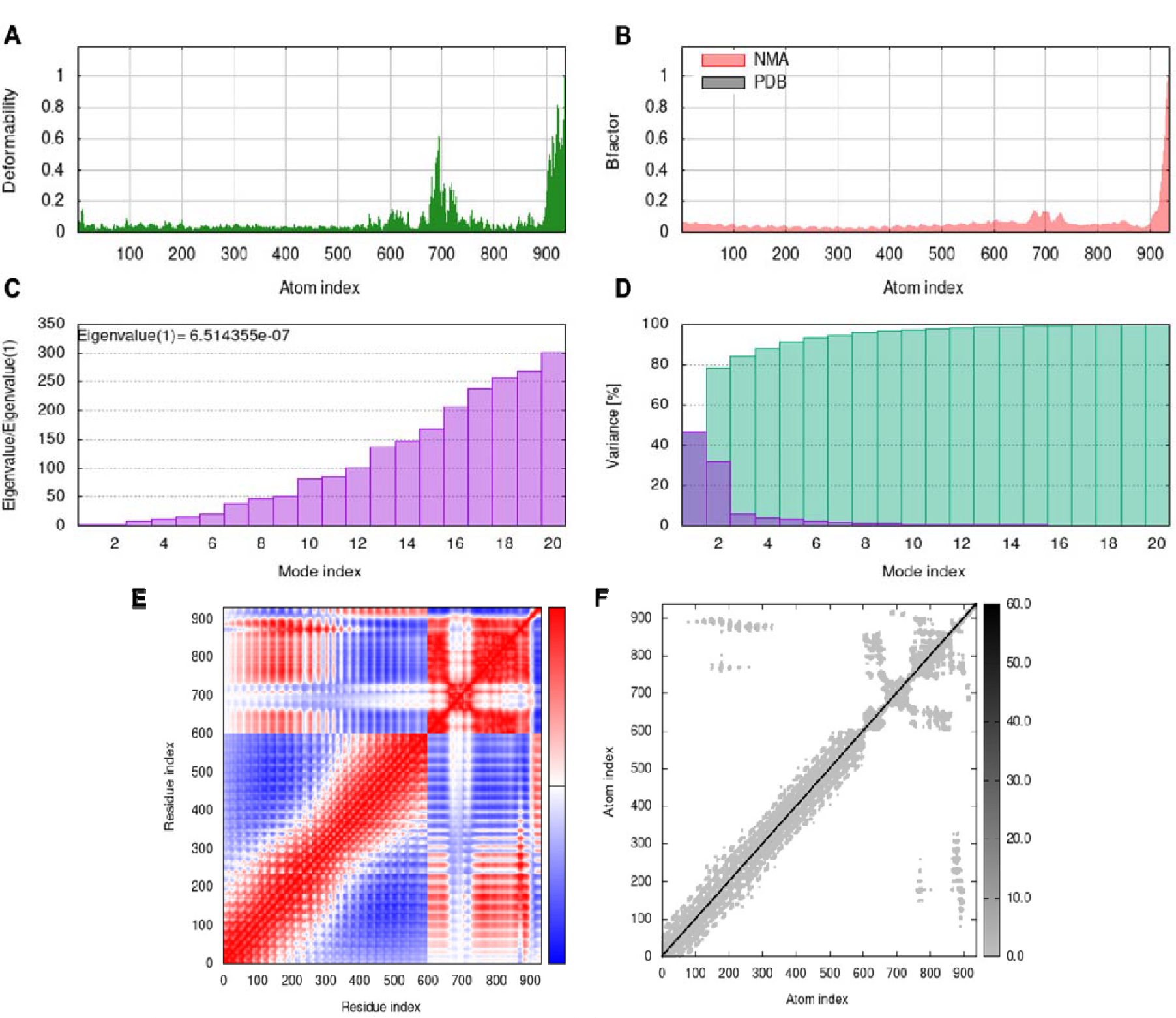
MD simulations of TLR4-vaccine complex by iMODS server. **(A)** Deformability, **(B)** B-factor analysis, **(C)** Eigenvalue, **(D)** Variance analysis, **(E)** Covariance matrix, **(F)** Elastic network map.

### Immune simulation

The C-IMMSIM immune server was utilized to simulate in silico immune responses with the candidate vaccine construct. Results after three doses of the vaccine showed an increase in secondary and tertiary immune reactions following the primary reaction. High concentrations of IgM and IgG demonstrated a humoral immune response (Figure 6A). The B cell population displayed an increase in B memory cells, which are important for prompt and effective recall responses, as well as an increase in B cell isotype IgM and a decrease in antigen concentration (Figure 6B, Supplementary Figure 1A). The TH1 and TC cell populations showed a significant response with associated memory development that raised antigen clearance on subsequent exposures (Supplementary Figure 1B-D). Increased numbers of dendritic cells that capture and process antigens for presentation by major histocompatibility complex (MHC) molecules to T cells and increased numbers of macrophages that excel in clearing invading pathogens and infected cells were observed (Supplementary Figure 1E-F). There were significantly higher numbers of natural killer cells that are important in innate immunity and fight against infected cells (Supplementary Figure 1G). In addition, the vaccine was able to produce a variety of cytokines, including IFN-gamma, IFN-beta, IL-10, and IL-23, which are crucial for inducing an effective immune response against pathogens (Figure 6C).

**Figure 6.**
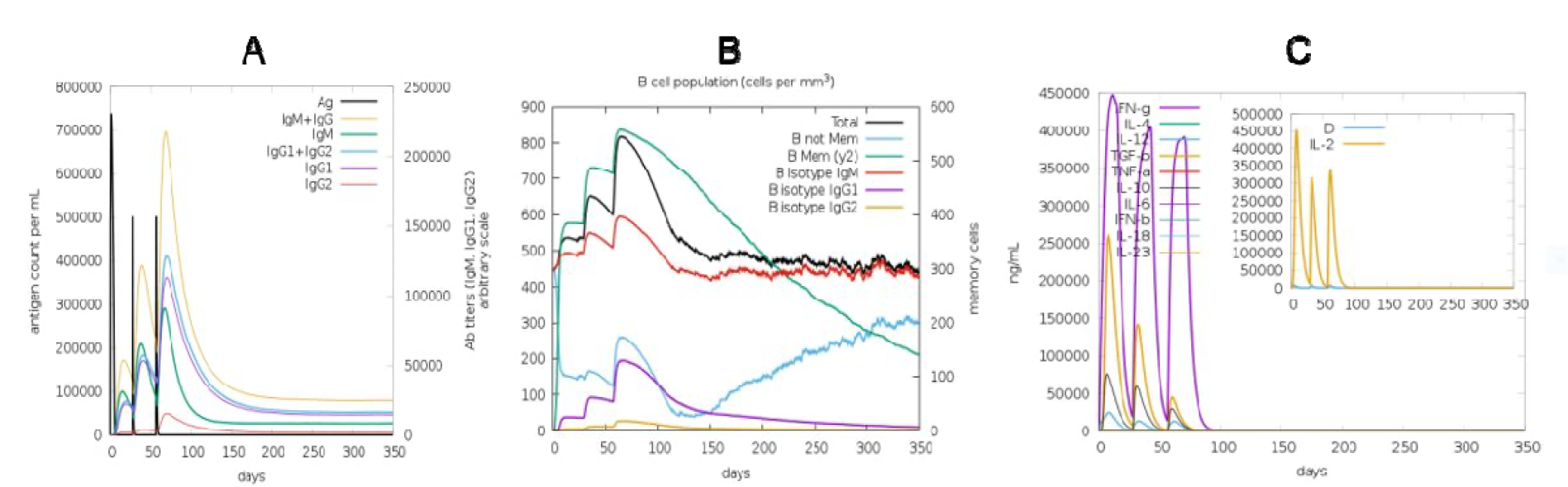
Using C-IMMSIM to simulate different immune responses of the vaccine construct: (**A)** Antigen and immunoglobulins, (**B)** B cell population, (**C)** obvious concentration of cytokines and interleukins.

### In silico cloning of the final construct

JCAT was utilized to optimize the codon usage of the engineered vaccine for a high level of expression in the *Escherichia coli* expression system. The vaccine construct was virtually integrated to facilitate optimal expression in the *E*.*coli* system. Codon production was executed, the designed vaccine showed a GC content of 52%, alongside CAI of 1.0, Subsequently, the recombinant plasmid was synthesized by incorporating the adapted codon sequences into the pET-28a (+) vector. XhoI and NedI sites were incorporated at the 5’ and 3’ termini of the vaccine sequence to enable the generation of sticky ends after restriction digestion. The total obtained length of circular pET28a(+) plasmid along with the insert was determined to be 6289 base pairs represented in Figure 7. Using SnapGene software, the process of visualizing and generating cloned maps was carried out.

**Figure 7.**
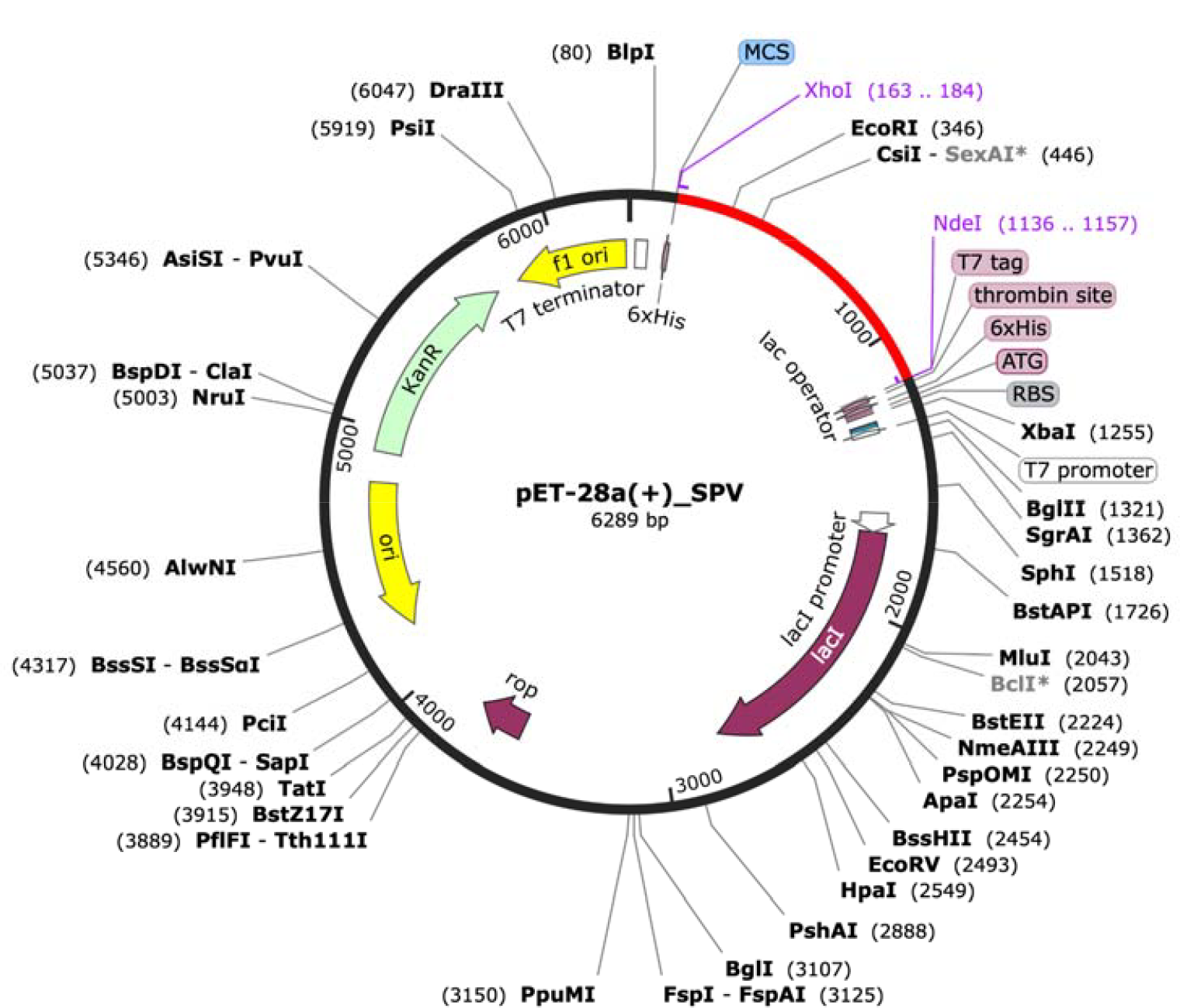
In silico cloning of the vaccine sequence into the pET28a (+) vector, represented in red color (6289 bp). The vaccine sequence was inserted between the XhoI and NdeI restriction sites of the expression vector.

## Discussion

Streptococcus pneumonia is a common respiratory pathogen that is associated with a high rate of morbidity and mortality among adults and children. Ranked as one of the highly virulent pathogens with a heavy disease burden, it can cause bacterial pneumonia, bacteremia, and meningitis (Tian, Zheng et al. 2023).Available SPN vaccines suffer from some limitations such as high production cost and low coverage which necessitates the desire to design a novel vaccine that combat pneumococcal infections. Using in-silico experiments, termed reverse vaccinology, it becomes possible to design an epitope-based vaccine which can be verified by in-vitro and in-vivo experiments. Our study aimed to design a multi-epitope vaccine that can elicit a strong immune response against SPN. DiiA protein was selected as a target for epitope prediction. It is a virulence factor that contributes to the long-term invasion and colonization of the pathogen. A previous study showed that immunization with DiiA reduced nasopharyngeal colonization and protected against invasive disease in mice.

DiiA was shown to be highly antigenic, non-allergenic and most importantly non homologous to human proteome, indicating its potential to be a candidate for vaccine construction. CTL, HTL and LBL epitopes were predicted and analyzed using various servers. Epitopes predicted to be highly antigenic, non-allergenic, nontoxic and can induce immune response were selected for vaccine design. Vaccine was constructed by combining CTL, HTL and LBL epitopes linked by AAY, GPGPG and KK linkers respectively. Linkers were added to minimize the junctional immunogenicity and to preserve each individual epitope identity during vaccine processing within the cells (Ayyagari, T et al. 2022). To boost the immune response, 50S ribosomal protein was attached to the vaccine as an adjuvant through the EAAAK linker. In-silico analysis of the constructed vaccine, composed of 332 amino acids, estimated that it would be highly antigenic and non-allergenic. In addition, it had the propensity to be highly stable, had high thermostability and was hydrophilic, predicted to be soluble upon overexpression. The vaccine was also estimated to have a long half lifetime. Molecular docking analysis between the vaccine and TLR4 showed very low binding energy, indicating the formation of a stable complex. Many favorable interactions (hydrogen bond, salt bridges and non-bonded interactions) contributed to this stable binding. To further confirm the stability of the docked vaccine, MD simulations were carried out using NMA. It showed rigid and stiff binding between the vaccine and the receptor. Ideal vaccine should be able to induce both cellular and humoral immune response. Immune simulation analysis showed that the designed vaccine has a significant level of immunoglobulins generation (IgM+IgG) following injection in addition to various interleukins and cytokines. Codon optimization showed satisfactory GC content and CAI value ensuring high expression of the vaccine in *E. coli* strain K12. The results of our study suggest the constructed vaccine as a potential candidate against SP infection. The antigenic epitopes predicted in this work can be used in future research to design novel multiepitope-based vaccines. However, the efficacy of the in-silico designed vaccine needs to be verified by wet lab experiments.

## Conclusions

Pneumococcal infection is a life-threatening disease affecting millions of people worldwide. It became of utmost importance to search for potential therapeutics for preventing this infection. Current vaccines suffer from several drawbacks which limit their efficacy. In this study, we investigated a novel protein target for vaccine design. Using several bioinformatics tools, we constructed a multi-epitope vaccine using DiiA protein as a target. Molecular docking, MD simulations, immune simulation and codon optimization results confirmed the construct as a potential vaccine candidate. High immunogenicity with no toxicity or allergenicity was also shown. So, the vaccine can be used as an alternative to conventional vaccines after validating its efficacy by in-vitro and in-vivo studies.

## Conflict of interests

The authors declare that no competing interests exist.

## Funding

No external funding was received.

## Notes

### Competing Interest Statement

The authors have declared no competing interest.

