## Supplementary file for "Multiepitope-based vaccine design against DiiA in *Streptococcus pneumoniae*, An immunoinformatics approach"

**Original article:**

**Supplementary Table 1. Physicochemical analysis of DiiA protein as predicted by ProtParam.**

| **Physicochemical property** | **Value** |
| --- | --- |
| Molecular Weight | 22648.16 |
| pI | 8.65 |
| Number of negatively charged residues | 26 |
| Number of positively charged residues | 28 |
| Estimated half-life | 30 hours (mammalian reticulocytes,in vitro) |
| Instability index | 47.54 (unstable) |
| Aliphatic index | 64.25 |
| Grand average of hydropathicity (GRAVY index) | -0.673 (hydrophilic) |

**Supplementary Table 2. Predicted HTL epitopes of DiiA protein predicted by NetMHCII-2.3 server.**

**Peptide**

ASLGLVVATSGFALL

LASLGLVVATSGFAL

ASSAALASLGLVVAT

SAALASLGLVVATSG

APKTTSAPQATAGTM

KTTSAPQATAGTMQD

PQATAGTMQDVTYQS

APNAAPKTTSAPQAT

VAPNAAPKTTSAPQA

QNSAQSQASKQLATE

TGSQNSAQSQASKQL

NSAQSQASKQLATEK

SAQSQASKQLATEKE

RPVAPNAAPKTTSAP

TILGKDTVQQSAKGE

PVAPNAAPKTTSAPQ

AGTMQDVTYQSPAGK

**Supplementary Table 3. Predicted linear B-cell epitopes of DiiA protein predicted by ABCpred.**

| **Rank** | **Sequence** | **Start position** | **Score** |
| --- | --- | --- | --- |
| 1 | SVTQEATPEYKLENTP | 17 | 0.96 |
| 2 | QDEIKGAPLSDKEKAE | 94 | 0.94 |
| 3 | GDKGGNTGSSDANANE | 34 | 0.93 |
| 4 | EYKLENTPGGDKGGNT | 25 | 0.91 |
| 5 | GGSQAGGSAHTGSQNS | 51 | 0.84 |
| 6 | PAGKQLPNTGSASSAA | 182 | 0.81 |
| 7 | NAAPKTTSAPQATAGT | 158 | 0.80 |
| 8 | EKAAKDKQDEIKGAPL | 87 | 0.79 |
| 8 | PKRPVAPNAAPKTTSA | 151 | 0.79 |
| 9 | TILGKDTVQQSAKGES | 2 | 0.78 |
| 9 | VQAIAMVTVPKRPVAP | 142 | 0.78 |
| 10 | ASKQLATEKESAKNAI | 71 | 0.77 |
| 10 | APQATAGTMQDVTYQS | 166 | 0.77 |
| 11 | MQDVTYQSPAGKQLPN | 174 | 0.76 |
| 11 | LKEIENAKTMEDVKEA | 122 | 0.76 |
| 11 | EAEKQAALKEIENAKT | 115 | 0.76 |
| 12 | KESAKNAIEKAAKDKQ | 79 | 0.74 |
| 13 | MEDVKEAETIGVQAIA | 131 | 0.71 |
| 14 | TGSQNSAQSQASKQLA | 61 | 0.68 |
| 14 | TGSSDANANEGGGSQA | 40 | 0.68 |
| 15 | TVQQSAKGESVTQEAT | 8 | 0.65 |
| 15 | SDKEKAELLARVEAEK | 103 | 0.65 |

**Supplementary Table 4. Predicted discontinuous B-cell epitopes.**

| **Residues** | **Number of residues** | **Score** |
| --- | --- | --- |
| A:L320, A:E321, A:N322, A:T323, A:P324, A:G325, A:G326, A:D327, A:K328, A:G329, A:G330, A:N331, A:T332 | 13 | 0.977 |
| A:T305, A:G306, A:S307, A:S308, A:D309, A:A310, A:N311, A:A312, A:N313, A:E314, A:K316, A:Y318 | 12 | 0.786 |
| A:K261, A:K262, A:T263, A:G264, A:S265, A:S266, A:D267, A:A268, A:N269, A:A270, A:N271, A:E272, A:G273, A:G274, A:G275, A:S276, A:Q277, A:A278, A:K279, A:G281, A:G282, A:S283, A:Q284, A:A285, A:G286, A:G287, A:S288, A:A289, A:H290, A:T291, A:G292, A:S293, A:Q294, A:N295, A:S296, A:K297 | 36 | 0.781 |
| A:F24, A:K27, A:E30 | 3 | 0.734 |
| A:K298, A:G299, A:D300, A:K301 | 4 | 0.672 |
| A:G302, A:G303, A:N304 | 3 | 0.672 |
| A:I66, A:L67, A:E68, A:A69, A:A70, A:G71, A:D72, A:K73, A:K74, A:G76, A:V77, A:I78, A:K79, A:V80, A:V81, A:R82, A:E83, A:S86, A:G87, A:L88, A:G89, A:L90, A:K91, A:E92, A:A93, A:K94, A:D95, A:L96, A:V97, A:D98, A:G99, A:A100, A:P101, A:K102, A:P103, A:L104, A:K117, A:A118, A:K119, A:L120, A:E121, A:A122, A:A123, A:G124, A:A125, A:T126, A:V127 | 47 | 0.656 |
| A:M1, A:A2, A:K3, A:L4, A:S5, A:T6, A:D7, A:E8, A:F28, A:E29, A:T31, A:F32, A:E33, A:V34, A:T35, A:A36, A:A37, A:V40, A:G221, A:P222, A:G223, A:P224, A:G225, A:K226, A:T227, A:S229, A:A230, A:P231, A:Q232, A:A233, A:T234, A:T237, A:M238, A:D240, A:G241, A:P242, A:G243, A:P244, A:G245, A:A246, A:P247, A:N248, A:A249, A:A250, A:P251, A:T253 | 46 | 0.655 |

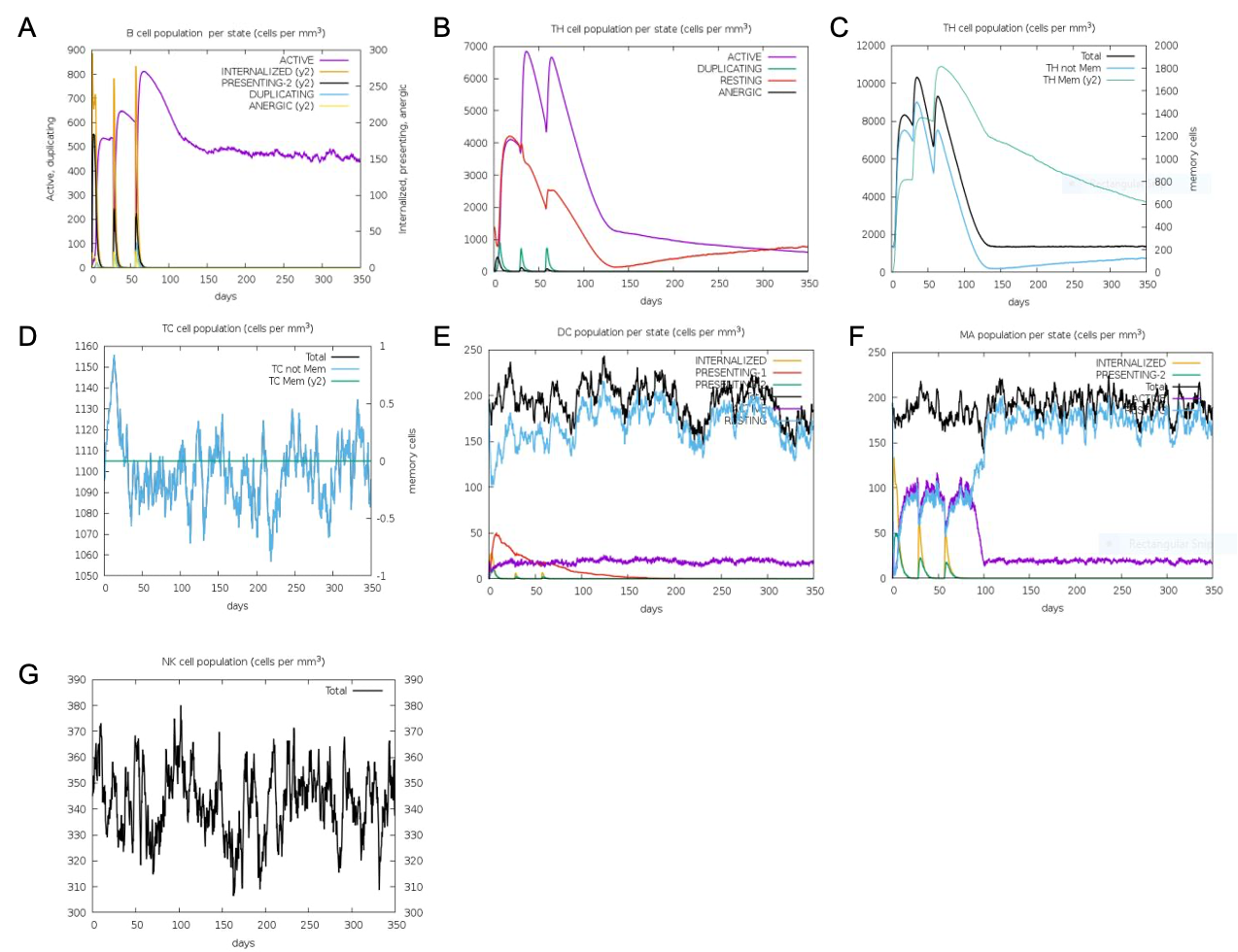

**Supplementary Figure 1. Using C-IMMSIM to stimulate different immune responses of the vaccine construct**: (**A)** B cell population per state, (**B)** helper T cell population, (**C)** helper T cell population per state, (**D)** cytotoxic T cell population per state, (**E)** dendritic cell population per state, (**F)** macrophage population per state, (**G)** natural killer cell population per state.
